# Transcriptional Landscape of THAP9 across Oligodendrocyte Developmental Stages

**DOI:** 10.1101/2025.01.17.633638

**Authors:** Tanuja Bhardwaj, Dhrumi Patel, Sharmistha Majumdar

## Abstract

Oligodendrocyte maturation and myelination are critical processes in human neurodevelopment, and their dysregulation is linked to numerous neurological disorders. While model organisms have provided insight into these processes, human-specific regulatory mechanisms remain poorly understood. This study investigated human THAP9, a protein homologous to the *Drosophila* P-element transposase, whose function in oligodendrocytes remains unknown. An analysis of publicly available RNA-sequencing data and H3K27ac ChIP-sequencing data from oligodendrocyte progenitor cells (OPCs) and mature oligodendrocytes (MOs) revealed significant upregulation of THAP9 during oligodendrocyte maturation. Co-expression analysis demonstrated a strong correlation with established markers of oligodendrocyte development, including myelin-associated genes (MOG, MBP) and key transcriptional regulators (PDGFRA, SOX5, SOX6, SOX11). THAP9 lacks homologues in mice, highlighting potential human-specific mechanisms in oligodendrocyte development and emphasising the importance of studying species-specific factors in neurodevelopment. Our findings suggest that THAP9 is a novel human-specific regulator of oligodendrocyte maturation and opens new avenues for studying myelination disorders.

## Introduction

Oligodendrocytes are glial cells in the central nervous system (CNS), responsible for forming the myelin sheath, a protective insulating layer surrounding and supporting axonal nerve fibers^1^. The mammalian CNS comprises approximately 70% myelin (by dry weight) ^2^, which acts as the metabolic, physiological and structural support of axons^3^. Degeneration or dysfunction of oligodendrocytes results in demyelination or dysmyelination, resulting in neurodegenerative diseases like multiple sclerosis^4^.

Understanding the maturation and development of oligodendrocytes is crucial for uncovering the regulatory mechanisms that govern these processes, providing valuable insights into neurodegenerative conditions. Neural stem cells give rise to OPCs during embryonic development^1^. Depending upon the brain region and the CNS developmental stage, oligodendrocyte precursor cells (OPCs), also called NG2 cells, can differentiate into oligodendrocytes, astrocytes and neurons^5^. However, the native fate of OPCs is to develop into oligodendrocytes^6^. The formation of MOs from OPCs occurs via many regulatory steps: OPCs differentiate into immature oligodendrocytes, which undergo various morphological changes to form mature and myelinating oligodendrocytes (MOs). This process is regulated by extrinsic factors, such as PDGF-α, IGF1, and Neuregulin-1, and intrinsic factors involving transcription factors, such as SOX10, OLIG1, and OLIG2 ^7^.

Transposons, or mobile genetic elements, comprise approximately 50% of the human genome^8^. Transposable elements (TEs) play important roles in neural stem cell differentiation into various neural lineages^9^. TEs act as sources of cis-regulatory elements, influencing gene regulatory networks, altering chromatin accessibility, and facilitating transcriptional regulation essential for the differentiation and function of these critical neural cells^10^. TEs can also integrate into regulatory networks that control myelination by providing additional regulatory elements that can enhance or suppress the expression of myelin-related genes, such as Myelin Basic Protein (MBP) and Proteolipid Protein (PLP)^11^. TEs have also been associated with various neurodegenerative conditions such as multiple sclerosis and dementia^12^; however, the molecular mechanisms through which TEs contribute to these conditions are not fully understood ^12^.

This study focuses on a domesticated DNA transposable element-derived gene called THAP9, a member of the THAP (THanatose-Associated Protein) family of proteins, which consists of 12 proteins (THAP1-THAP12)^13^ in humans. THAP9 gene maps to chromosome 4q21.22 and consists of 7 exons, though the exact numbers can vary depending on splice variants^14^. The THAP9 protein is a *Drosophila* P-element transposase (DmTNP) homologue^15^. The members of the THAP family are involved in functions including apoptosis^16^, cell proliferation^16^, pluripotency^17,18^ and differentiation of embryonic stem cells (ESCs)^19^. According to the Human Protein Atlas ^20^, THAP9 is expressed in the brain, and single-cell studies suggest its enriched expression in oligodendrocytes. However, its specific role in oligodendrocytes remains unknown,

Most studies on oligodendrocyte development have been performed in model organisms such as mouse^21^. However, certain differences exist in the biology of oligodendrocytes of humans and rodents ^21^, resulting in a lacuna of knowledge about human-specific mechanisms that result in oligodendrocyte maturation and myelination. We hypothesised that since THAP9, which is highly expressed in oligodendrocytes, has no homologues in mice; it could be potentially important for human-specific oligodendrocyte functions or maturation.

In this study, we investigated the role of the THAP9 gene in oligodendrocyte biology by studying its regulation during the maturation and differentiation of OPCs into MOs. Our results suggest that THAP9 could potentially be involved in oligodendrocyte lineage-specific development and myelination.

## Results

### THAP9 Shows Consistent Upregulation During Human Oligodendrocyte Maturation Across Developmental Stages

The process of oligodendrocyte maturation is important for proper myelination in the human brain; despite the importance of this vital process, human-specific aspects of this process remain poorly understood. We analysed RNAseq data (submitted in GEO ^22,23^) of oligodendrocyte precursor cells (OPCs) and mature oligodendrocytes (MOs) isolated from the gray matter of the dorsolateral prefrontal cortex of human donors (10 infants samples aged 0-2 years; Figure 1:Ia) and 12 adults samples aged 30-40 years, Figure 1:IIa). Each cell type forms a distinct cluster indicating different expression profiles (Figure 1:Ia, IIa). Differential expression analysis revealed extensive transcriptional changes during oligodendrocyte maturation in both age groups. In infant samples (Figure 1:Ib), we identified 17,197 differentially expressed genes (9,047 downregulated, 8,150 upregulated). Adult samples (Figure 1: IIb) showed similar numbers with 19,430 differentially expressed genes (10,479 downregulated, 8,951 upregulated). THAP9 showed consistent and substantial upregulation in MOs compared to OPCs in both age groups [Figure 1: Ic (Infants: log2 fold change of 1.46 (MOs vs OPCs) with an adjusted p-value of 7.117729e-12) and IIc (Adults: log2 fold change of 1.34 (MOs vs OPCs) with an adjusted p-value of 1.17208e-47)].

**Figure 1:**
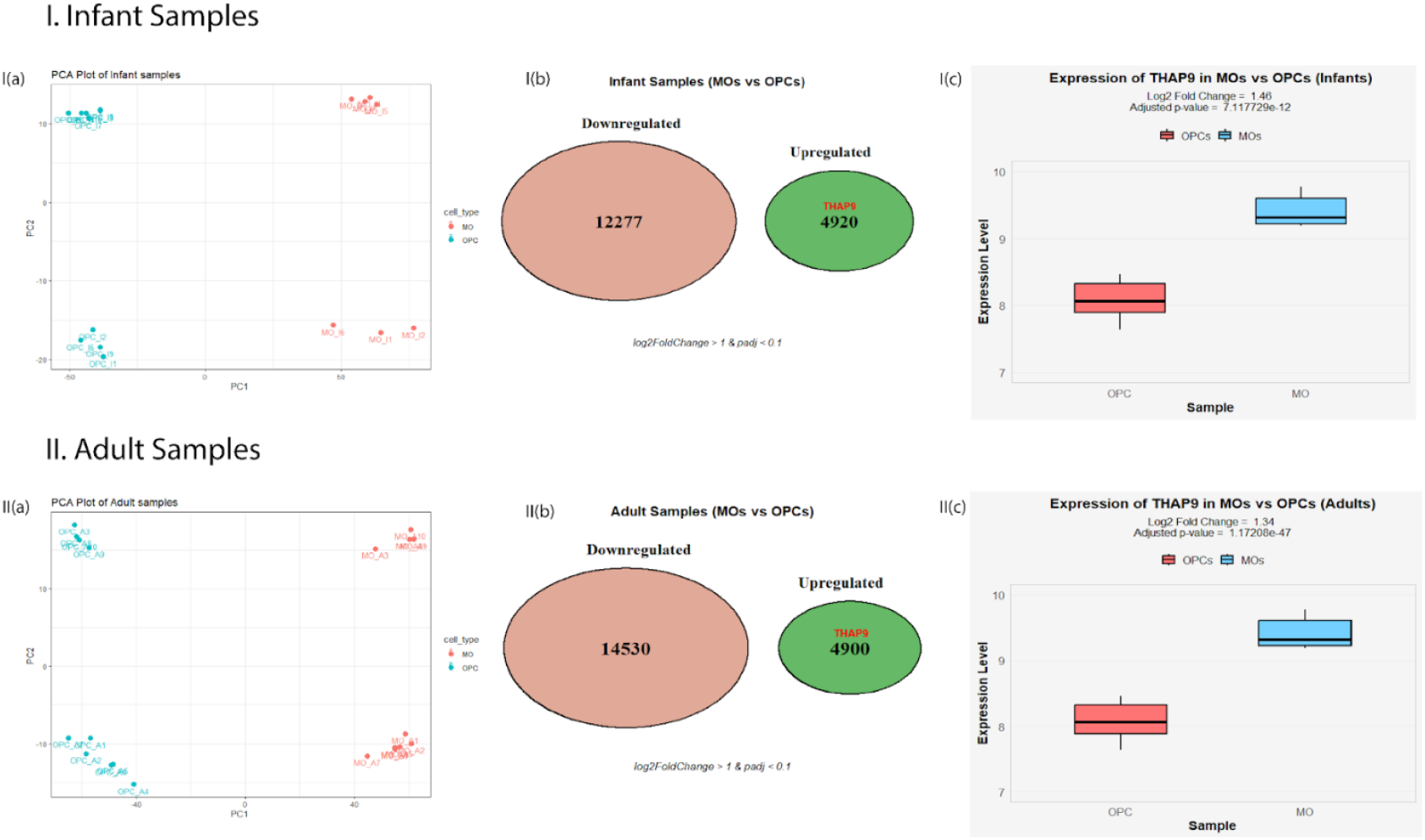
Transcriptional Analysis of Human Oligodendrocyte Maturation in Infant and Adult Samples: (A) *PCA plot* of I.Infant and II. Adult samples. MOs (red dots), OPCs (turquoise dots). (B) *Venn diagram showing differential expression analysis comparing MOs and OPCs*, I. In infant samples; Downregulated genes (salmon-coloured circle;12277), Upregulated genes (green-coloured circle; 4920 genes, including THAP9) II. In Adult samples; Downregulated genes (14530); Upregulated genes (4900 genes, including THAP9) [Statistical criteria: log2FoldChange > 1, Adjusted p-value (p-adj) < 0.1 (indicating statistical significance with FDR correction)]. (C) *Boxplot representing THAP9 gene expression levels* in MOs (blue box) and OPCs (red box) in I. infant II. adult samples. Expression levels are shown on a log2 scale (y-axis). Statistical analysis was performed using DESeq2.

Thus, through transcriptomic analysis of human dorsolateral prefrontal cortex samples, we identified THAP9 as a consistently upregulated gene during oligodendrocyte maturation in infant and adult brain tissue (Suppl. Fig. 1). It is to be noted that most studies about oligodendrocyte maturation and myelination are performed in mice models. Interestingly, THAP9 lacks a homolog in rodents like mice and rats, suggesting its potential human-specific role in myelination.

### Epigenetic signatures suggest that THAP9 is transcriptionally more active in MOs compared to OPCs

We next decided to investigate if the higher expression of THAP9 in MOs than in OPCs could be explained by its altered epigenetic regulation in these cell types. H3K27ac (histone H3 lysine 27 acetylation) is a histone modification associated with active enhancers and promoters and is associated with positive transcriptional activity of a downstream gene. The THAP9 gene is located on the forward strand of chromosome 4, at position 82,900,684-82,919,969 (GRCh38). H3K27ac ChIP-seq data revealed a conserved epigenetic signature at this locus, wherein enriched peaks (Figure 2c) were observed in both infant and adult MOs, suggesting that THAP9 is transcriptionally more active in MOs than in OPCs (Figure 2 a,b). Although there is a slight increase in the H3K27ac peak signal in OPCs from infants to adult (Figure 2 a,b), indicating some regulatory activity in these cells, the enrichment of H3K27ac in MOs indicates a cell-type-specific regulation of THAP9. The stable H3K27ac signal in MOs across developmental stages implies that THAP9 plays a consistent regulatory role in mature oligodendrocytes, while its activity in OPCs is comparatively lower. These results are consistent with our observation that THAP9 is upregulated (in MOs compared to OPCs) and suggest that the gene plays a role during oligodendrocyte differentiation.

**Figure 2:**
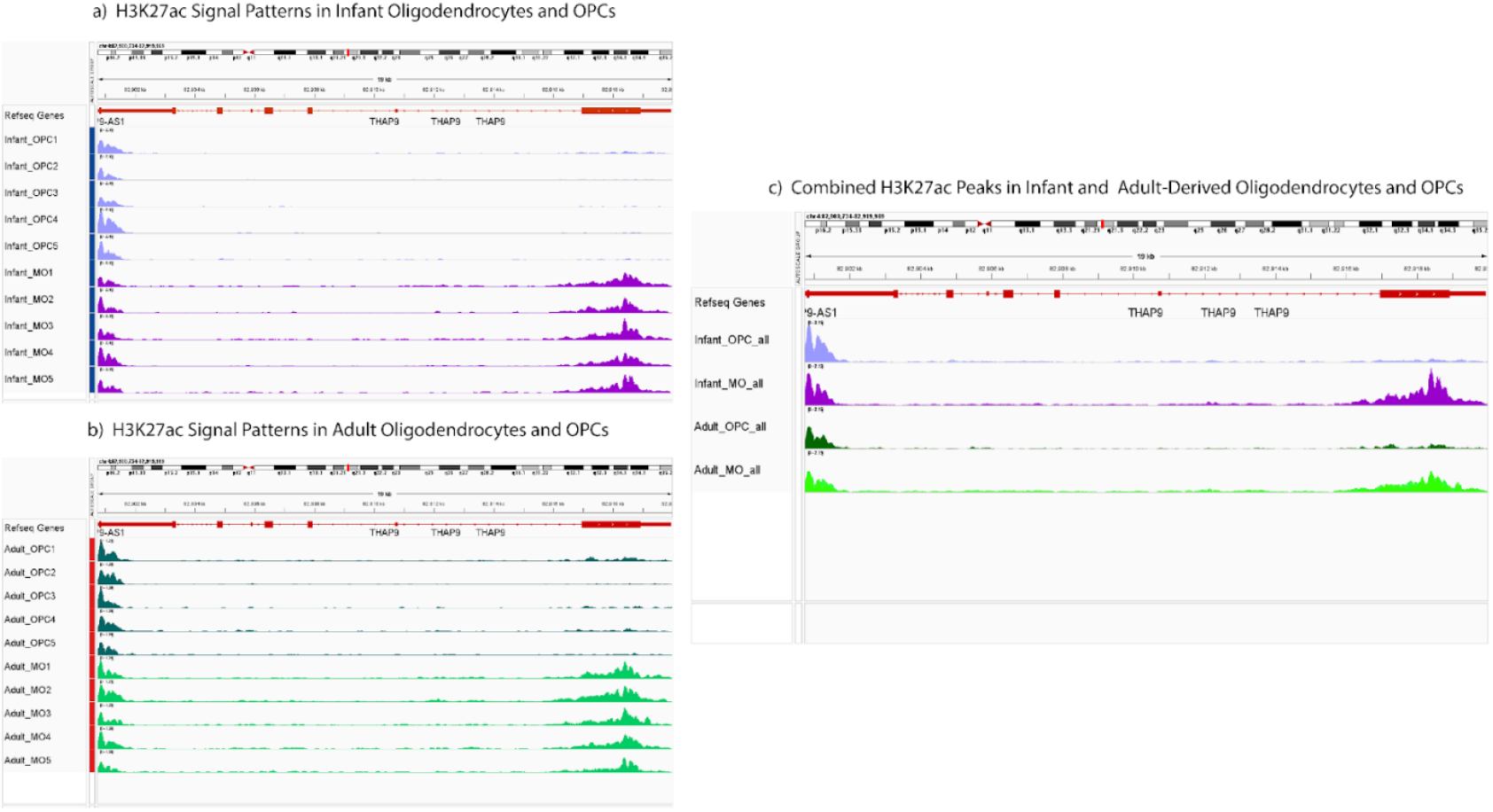
Histone H3K27ac enrichment at the THAP9 gene locus in OPCs and MOs from adult and infant brain samples. H3K27ac ChIP-seq track plots across the THAP9 gene for (a) infant samples for OPC (lavender) and MO (purple). (b) adult samples for OPCs (dark green) and MOs (light green) (c) Combined H3K27ac ChIP-seq track plots across the THAP9 gene for infant MO (purple peaks), adult MO (light green peaks), adult OPC (green peaks), infant OPC (purple peaks). Individual H3K27ac ChIP-seq tracks in THAP9 locus

### Transcriptional Regulation, not alternative splicing, causes changes in THAP9 expression in MOs versus OPCs

To investigate the possible regulatory mechanisms that could dictate THAP9 expression during cellular development and maturation, we examined the expression of its alternate isoforms. Isoform switching or the differential usage of alternatively spliced isoforms of a gene can control its function, localisation or stability of the protein it encodes. Both expression and splicing of a gene can change in a spatiotemporal fashion and often vary during stages of development.

Thus, the isoform switching of the THAP9 gene was investigated in both MOs and OPCs across infant and adult samples to identify if the gene is regulated during developmental transitions in these cell types.

The results obtained included information about gene expression, isoform expression and isoform usage. Gene expression represents the measure of total transcriptional output of the whole THAP9 gene, which is calculated by the sum of expression of all its isoforms (in TPM, Transcript Per Million); Isoform expression measures the quantity of a specific isoform individually by calculating the normalised read counts for each THAP9 transcript variant (labelled with unique ENST number); Isoform usage (isoform fraction - IF) measures the relative amount of each isoform compared to the total gene expression. Isoform switching refers to a condition when there is a statistically significant shift in preferential usage of a specific isoform.

The isoform switch analysis demonstrated that the THAP9 gene did not undergo significant change in isoform usage or switching in MOs and OPCs in samples obtained from adults and infants (Figure 3). Moreover, the gene shows consistently higher expression of all its isoforms in MOs (Figure 3 a, b bottom left and middle panels) as compared to OPCs. Thus, THAP9 regulation appears to be primarily driven by transcriptional control rather than alternative splicing with conserved expression patterns of the gene in adults as well as infant cells. Among all the isoforms, ENST00000506208 (highest in MO_Infant) and ENST00000514440 (highest in MO_Adult) show high expression levels in both adults and infants. In infants (Figure 3a), ENST00000514440 appears to have higher usage (∼ 0.3 or 30%, right panel) in OPCs; however, statistical analysis suggests that this difference is insignificant and cannot be considered an “isoform switch”. Similarly, in adult samples (Figure 3b), although ENST00000514244 has lower expression in OPCs (middle panel), no significant switch between MOs and OPCs (right panel) was observed. Thus, we concluded that in both infants and adults, there was no significant change in THAP9 isoform usage in either cell type. These results show that THAP9 maintains its cell specificity with conserved regulatory mechanisms in both stages of life.

**Figure 3:**
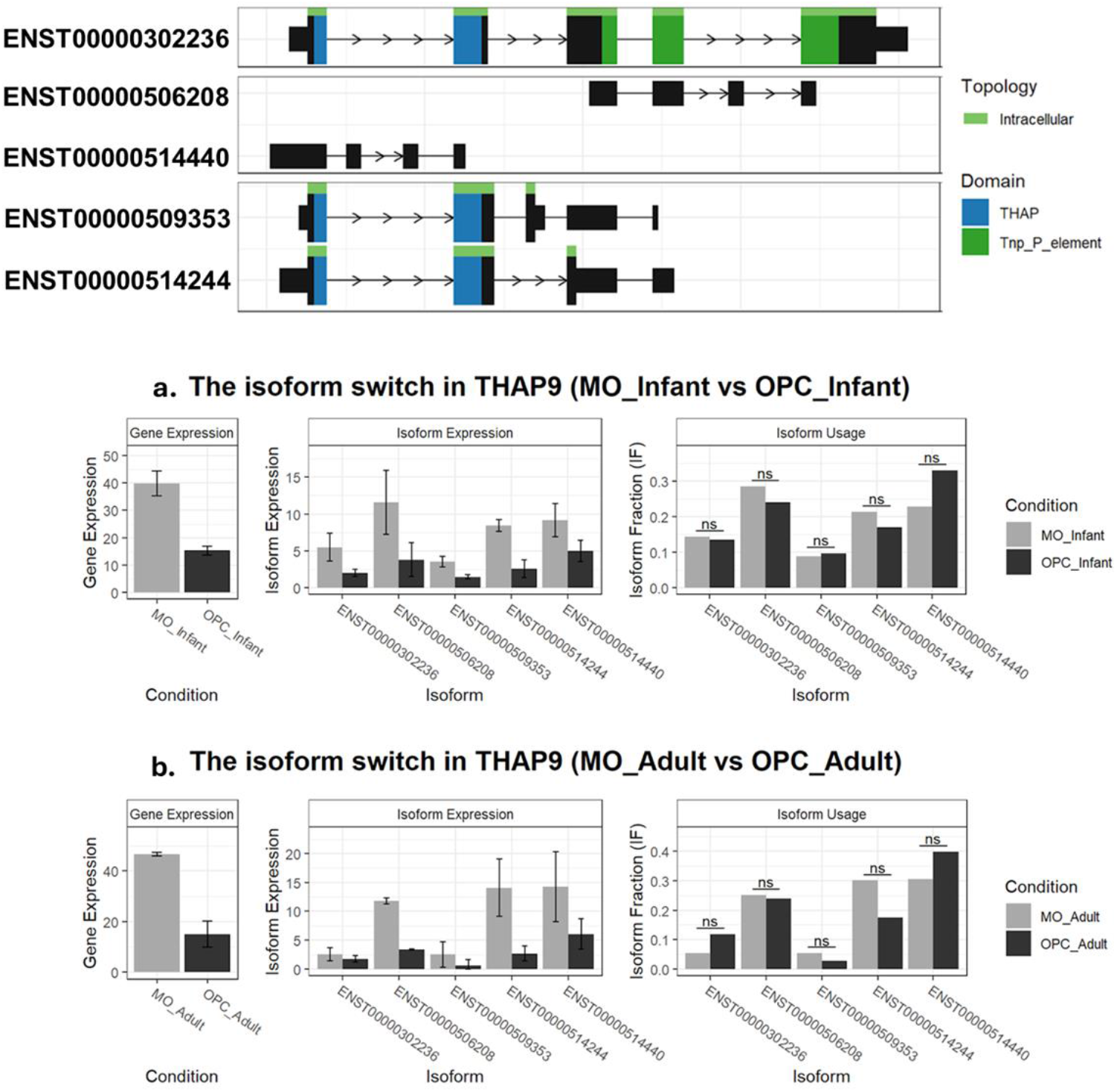
Isoform switch analysis of THAP9 in Mature Oligodendrocytes (MO) and Oligodendrocyte Progenitor Cells (OPC). *Top panel*. Five different THAP9 isoforms with ENST numbers, THAP domain (blue) and Tnp_P_element domain (green) **(a)** In MO_Infants: Gene Expression (left panel) represents overall THAP9 gene expression; Isoform Expression (middle panel), represents the amount of specific isoform expressed; Isoform Usage (right panel) represents relative expression of an isoform **(b)** In MO_Adults: Gene Expression (left panel); Isoform Expression (middle panel); Isoform Usage (right panel).

### Gene co-expression analysis of THAP9 suggests its potential role in oligodendrocyte development

To understand the function and regulatory mechanisms of THAP9 in oligodendrocyte cell types, we performed weighted gene co-expression analysis (WGCNA). WGCNA helps in the identification of a co-expression module in which a particular gene of interest lies, thus providing insights into its associations with different cellular states (e.g., OPCs or MOs across developmental stages). By analysing these modules, WGCNA allows us to understand the possible regulatory and biological networks in which a gene functions. In this study, WGCNA was performed to assess THAP9 co-expression patterns between OPCs and MOs, elucidating the gene’s system-dependent roles in cellular differentiation and maturation processes. We performed WGCNA separately for infant and adult samples of MOs and OPCs for network construction with two datasets-one containing only protein-coding genes (Suppl. Fig. 2 for infant samples, Suppl. Fig. 3 for adult samples) and one containing all genes (comprehensive analysis, Figure 4). We then performed hierarchical clustering analysis on infant and adult samples, showing distinct clustering patterns between the two groups. Infant samples were categorised as MOs (green) and OPCs (orange) (Suppl. Figure 4a). Similarly, adult samples were grouped into OPCs (green) and MOs (orange) (Suppl. Figure 4d) and block dendrograms displaying block sizes, with infant samples containing 8893 genes in block 1 and 5597 genes in block 2 (Suppl. Fig. 4b,c) and adult samples containing 8558 genes in block 1 and 5875 genes in block 2 (Suppl. Fig. 4e,f), which included both coding and non-coding genes.

**Figure 4:**
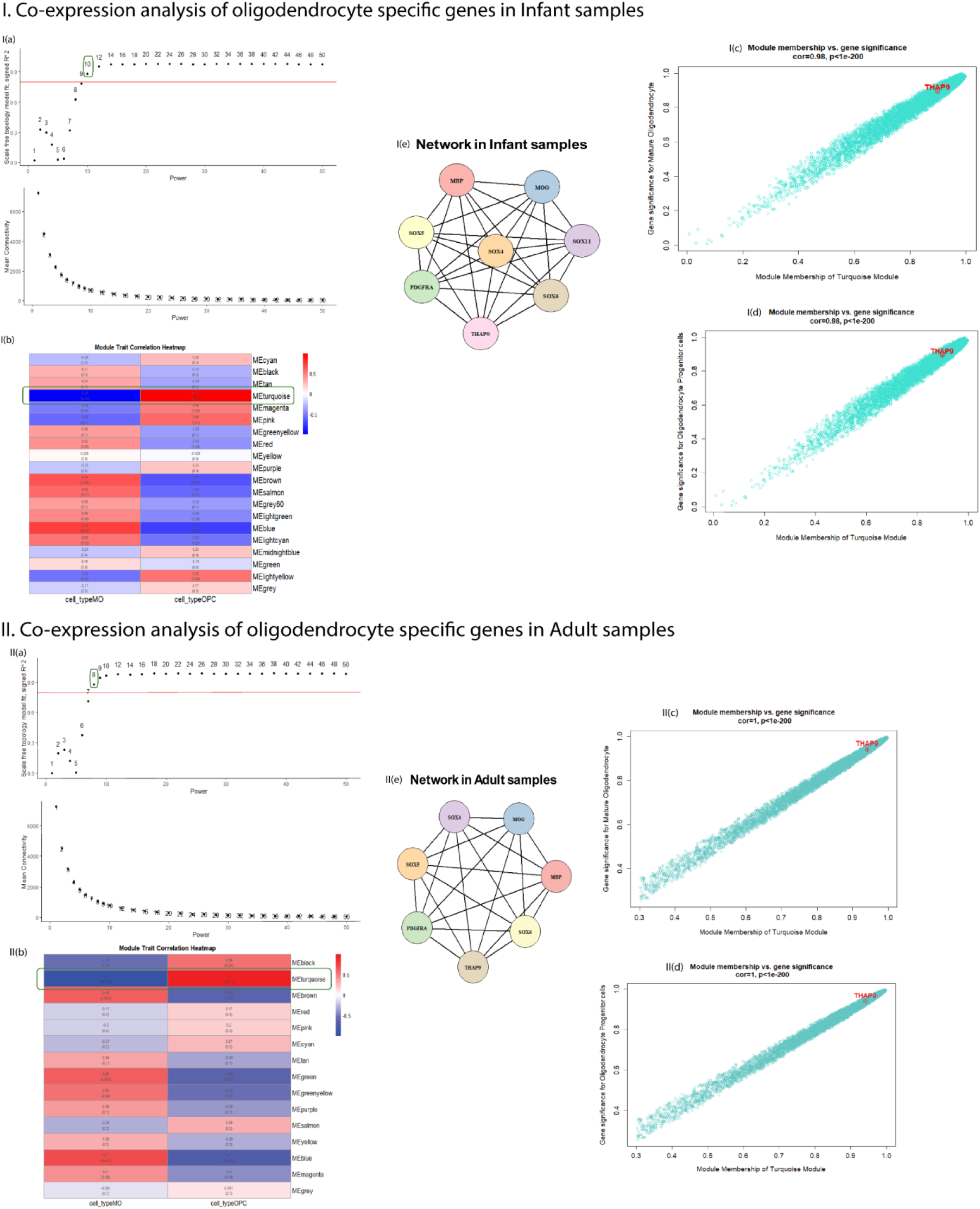
Co-expression network analysis of THAP9 and associated genes obtained by comprehensive analysis using WGCNA. **I(a)+II(a)** *Scale mean independence and mean connectivity graphs*. The top graph represents Scale-free topology model fit (R^2^, Y-axis) vs soft-threshold power (X-axis). The red line indicates the R^2^ = 0.8 threshold, and the numbers indicate the different powers tested. The lower graph represents the mean connectivity (Y-axis) vs soft-thresholding power (X-axis) and shows a decrease in connectivity as the power increases. Selected soft-thresholding power (ß, circled in green) = 10 (infants) and 8 (Adults). **I(b)+II(b)** *Module-Trait Correlation Heatmap* displays the correlation between module eigengenes (ME) and two cell types (MOs and OPCs) in Infant (I) and adult (II) samples. The turquoise module (represented as MEturquoise) contained the THAP9 gene. Positive correlation (red), negative correlation (blue), corresponding correlation values and p-values in parentheses. **I+II(c**,**d)** *Module Membership vs Gene Significance* (OPCs and MOs) shows a strong positive correlation in the infant (I: cor=0.98, p<1e-200) and adult (II:cor=1, p<1e-200) samples, between module membership (MM) and gene significance (GS) for the Turquoise module. The scatter plots reveal a strong linear relationship, with genes showing higher MM with greater GS. The dense clustering of points along the diagonal indicates highly coordinated gene expression within the Turquoise module. THAP9 (marked in red) appears at the right corner in both plots, showing high MM (∼0.9) and GS (∼0.9) in both MOs and OPCs, indicating that THAP9 is a highly connected “hub gene” within the turquoise module in both infant (I) and adult (II) samples. **I(e)+ II(e)** *Co-expression network* derived from WGCNA demonstrates consistent co-expression patterns between THAP9 and oligodendrocyte regulatory genes (SOX4, SOX5, SOX6, SOX11) and marker genes (MBP, MOG and PDGFRA) in both MOs and OPCs in infant (I) and adult (II) samples. Each node represents a gene, and edges indicate a co-expression relationship.

In comprehensive (Figure 4 Ib + IIb) and coding gene-specific analysis (Suppl. Fig 2e, 3e) across adult and infant samples, the THAP9 gene was consistently found in the turquoise module (represented as MEturquoise) showing significant Module trait relationship of 0.98 (infants, Suppl. Fig.3, Fig 4) and 1.0 (adults, Suppl. Fig.3, Fig 4)] but opposing correlations with the two cell types (MOs and OPCs), suggesting the cell-type-specific expression pattern of the genes in this module. Turquoise module genes are predicted to be interconnected with high edges (<u>Supplementary file 2</u>) in a biologically important gene network in oligodendrocyte cells since they show a strong positive correlation between module membership and gene significance (Figure 4: I+II(c,d); Suppl. Fig 3f+g, 4f+g). When considering all genes, the infant samples contained slightly fewer genes and edges than the adult samples (<u>Supplementary file 2c + 2d</u>). When the analysis was restricted to coding genes, the turquoise module (Suppl. Fig. 2e+3e) in infant samples, unlike the corresponding adult samples, showed further reduction in the number of genes and edges (<u>Supplementary file 2a + 2b</u>).

To further refine our network, we extracted the THAP9 gene from the edge file in the turquoise module. We filtered to include only genes of interest (i.e., the earlier-mentioned oligodendrocyte regulatory and marker genes), thereby constructing a focused gene interaction network (Figure 4: Ie+ IIe). In both MOs and OPCs, THAP9 co-expressed (Figure 4: Ie+ IIe; Suppl. Fig 3h, 4h) with well-known oligodendrocyte lineage markers like MBP (Myelin Basic Protein, important myelin structural protein in MOs^24^), MOG (Myelin Oligodendrocyte Glycoprotein, key maturation marker and competent of myelin^25^), PDGFRA (Platelet Derived Growth Factor Receptor Alpha, important for OPC proliferation and maturation, important marker for oligodendrocyte precursor cells^26^), SOX5 and SOX6 (essential transcription factors for oligodendrocyte development^27^) and SOX4 and SOX11 (involved in early OPC specification^28^). All these genes had high-weight connections (weight>0.4, <u>Supplementary file 2a + 2b</u>) in coding genes. THAP9 co-expression patterns with precursor (PDGFRA) and mature (MBP and MOG) markers gave us insights into the gene’s involvement in multiple stages of oligodendrocyte differentiation. The co-expression with SOX genes indicates that THAP9 may play a role in the transition from OPCs to MOs through its role in differentiation and myelination processes. It is to be noted that THAP9 has a similar co-expression network architecture in adult and infant samples. Thus, THAP9 could be a part of an unrecognised conserved regulatory network controlling oligodendrocyte lineage progression and maintenance of myelination in human-specific oligodendrocytes.

Interestingly, while the important oligodendrocyte-related genes (MOG, MBP, SOX4, SOX5, SOX6 and PDGFRA) were all present within the turquoise module in both age groups, we observed some differences in their strength of correlations (<u>Supplementary file 2c + 2d</u>). In adult samples, MOG, MBP, SOX4, SOX5, SOX6 and PGDFRA maintained a strong correlation (weight >0.4, <u>Supplementary file 2d)</u> with THAP9. However, in infant samples, MOG and MBP had lower correlation values (weight ∼0.2, <u>Supplementary file 2c</u>) with THAP9, while SOX4, SOX5, SOX6 and PDGFRA retained high correlation values (weight ≥0.4, <u>Supplementary file 2c</u>). This difference demonstrates that including non-coding genes increases the number of genes and the number of connections (edges) within the turquoise module. Non-coding genes contribute additional layers of regulatory complexity and network connectivity.

This comprehensive analysis of coding and non-coding genes resulted in a more robust understanding of the transcriptional landscape and provided novel insights into the possible involvement of THAP9 in oligodendrocyte development. This is exemplified by THAP9’s consistent module placement and high trait relationship (Suppl. Fig. 3,4; Figure 4). However, the variations observed between the comprehensive and coding gene analysis highlight the complex molecular landscape of oligodendrocyte development biology. For instance, the comprehensive analysis gives a more detailed network pattern (<u>Supplementary file 2c + 2d</u>). On the other hand, the coding gene-specific analysis demonstrates a network wherein THAP9 shows more conserved strong co-expression with specific developmental genes in infant samples (Figure 4: Ie, Suppl. Fig. 2h). Conversely, adult samples display slightly different molecular interactions (Figure 4: IIe, Suppl. Fig. 3h), with variations in the gene co-expression relationships (i.e., topological overlap measures between genes) and absence of some genes such as SOX11 (weight < 0.2) (<u>Supplementary file 2d</u>). These patterns highlight the characteristics of ongoing developmental processes and early glial and neural maturation. The high module trait relationship (Fig I+II c,d) suggests that while the fundamental molecular mechanism remains the same, some subtle regulatory changes exist across the developmental stages. Despite these variations, the overall network structure and the strength of gene interactions remained consistent across conditions, whether analysing all genes or only coding genes. This consistency emphasises the robustness of the transcriptional networks governing oligodendrocyte development.

### THAP9 expression during direct reprogramming of human fibroblasts to Oligodendrocyte lineage cells

To understand the role of THAP9 in oligodendrocyte lineage cells, we further studied its expression in mature cells that have been reprogrammed to oligodendrocyte lineage. Reprogramming a mature cell into an oligodendrocyte lineage requires re-establishing all the epigenetic states and cell identity; this can provide insights into a gene’s role in lineage-specific transcription or the pathways essential for transition and oligodendrocyte lineage specification. Detailed expression analysis can help us understand whether a gene is required in earlier cell commitment or for terminal differentiation and myelination.

Towards this, we analysed RNAseq data^29^ from a study in which oligodendrocyte lineage cells were obtained by reprogramming human fibroblasts using a small molecule cocktail (A83-01, thiazovivin, purmorphamine, VPA, and forskolin) and OCT4 as induction factor. The study used two reporter systems as indicators for reprogramming success to track and isolate cells at specific developmental stages: OLIG2+ (OLIG2::eGFP) cells marking early stages and SOX10+ cells (SOX10::eGFP) to mark the progression to more mature stages. Different stages of the cells were made using specific markers: OLIG2, SOX10, O4+ and MBP (Suppl. Table1).

Figure 5d shows that in reprogrammed cells, THAP9 is highly expressed in early reprogramming stages of induced fibroblasts [A2B5(DF), A2B5(BJ)] but decreases in later stages of induced OPCs[O4(D40), O4(D60)], indicating a potential role in initiating lineage conversion. THAP9 thus displays distinct expression patterns in native (Figure 1: Ic+IIc) versus reprogrammed cells (OPCs derived from induced pluripotent stem cells), suggesting its role may differ between natural development and induced reprogramming where it is upregulated in MOs when compared to OPCs. As OPCs mature, although THAP9 expression decreases, it remains elevated in cells expressing key transcription factors involved in oligodendrocyte differentiation, such as OLIG2 and SOX10. This suggests a functional relationship between THAP9 and OLIG2 (Figure 5a, d).

**Figure 5:**
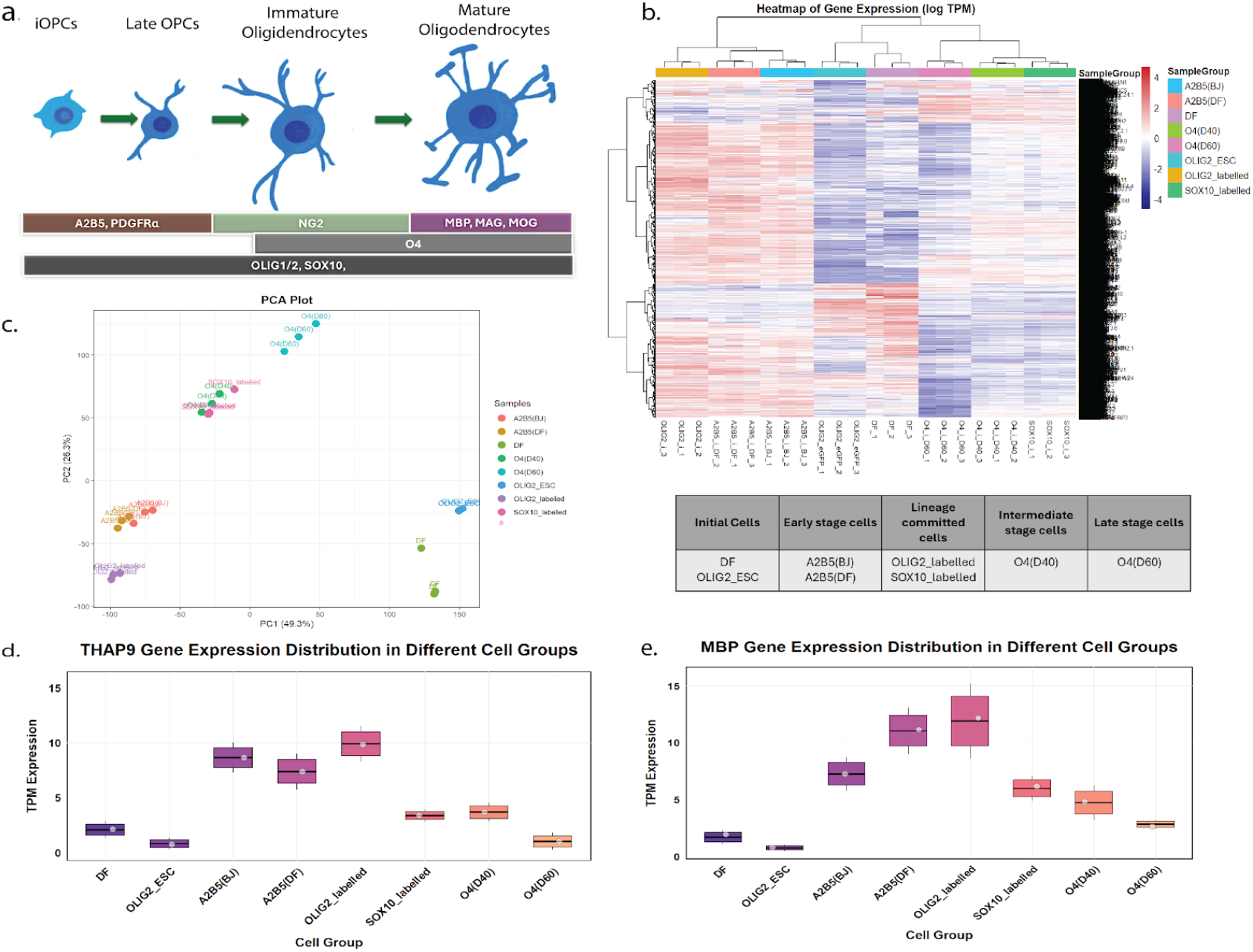
Gene expression analysis in fibroblasts reprogrammed into maturing oligodendrocytes shows different transcriptomics profile. **a)** *Specific marker proteins associated with each stage of oligodendrocyte development*. Markers for the progression from immature oligodendrocyte precursor cells (iOPCs) to late OPCs, immature oligodendrocytes, and mature oligodendrocytes. **(b)** *Heatmap of gene expression in logTPM*. High expression (red), low expression (blue), intermediate (white) expression. Rows represent different genes; columns represent different samples with colour-coded legends (on the right). The top of the dendrogram shows the hierarchical clustering of the samples. (**c)** *PCA (Principal Component Analysis) plot* shows the relationship between the samples in two dimensions, PC1 (x-axis) and PC2 (y-axis), with each coloured dot representing a sample. The samples clustered together show similar gene expression patterns, and the samples with different expressions are in separate groups. **(d+e)** *Expression analysis of THAP9 (d) and MBP (e) across diverse cell populations*. Box plots depict the distribution of gene expression levels (d: THAP9, e: MBP) measured in Transcripts Per Million (TPM) among eight distinct cell groups; the boxes represent interquartile ranges with median values indicated by horizontal lines, while whiskers extend to the minimum and maximum values excluding outliers. Grey dots represent mean expression values for each group

Similar expression patterns were observed for the MBP (Myelin Basic Protein) gene, which is essential for the formation and functions of MOs^30^. MBP, like THAP9, has higher expression in the early stages of oligodendrocyte development (A2B5+, OLIG2+ cells), which decreases through maturation; this could signify that both genes are co-regulated during reprogramming. Moreover, both genes show similar expression patterns (Figure 5d+e) to OCT4-mediated reprogramming. This is corroborated by our co-expression analysis which also demonstrated that THAP9 co-expresses with oligodendrocyte development-specific genes, such as MOG and MBP.

MBP usually has high expression in mature cells, indicating reprogramming may need more optimisation to replicate natural oligodendrocyte development. It also signifies that O4(D60) cells have undergone incomplete maturation (Figure 5e). This analysis suggests that the oligodendrocytes obtained by reprogramming human fibroblasts do not completely mimic their natural developmental trajectory since there are marked differences in their transcriptomes and the time durations and morphologies required to express relevant markers.

Overall, the observed expression patterns suggest that THAP9 plays prominent roles in both the early stages of oligodendrocyte progenitor cell development and in regulating oligodendrocyte development and differentiation. Further functional studies would be necessary to understand the precise mechanisms by which THAP9 contributes to these processes.

## Discussion

Oligodendrocytes are cells responsible for myelination. Oligodendrocyte progenitor cells mature and form mature myelinating oligodendrocytes. Oligodendrocytes from different developmental stages exhibit different myelination abilities^31^. The proliferation, maturation and maintenance of oligodendrocytes is important since dysregulation in these processes results in conditions like neurodegenerative diseases or delayed repair ability after injury^32^.

In this comprehensive bioinformatics investigation, we studied the role of THAP9 in oligodendrocyte development. THAP9, encoded by a DNA transposable element-derived gene, is homologous to the P-element transposase of *Drosophila melanogaster* and is a member of the human THAP protein family of 12 members ^33,34^. Our analysis revealed a pattern of THAP9 upregulation during oligodendrocyte maturation, consistently observed across infant and adult tissue samples (Figure 1: Ic+IIc in infant and adult samples; 1.46 and 1.34 log2FoldChange, respectively). The analysis of histone modification H3K27ac across the THAP9 gene locus in MOs and OPCs provided us with further confirmation that THAP9 indeed shows more transcriptional activity in the MOs than OPCs. This persistent and stage-specific expression profile highlights that the THAP9 gene may play an important role in oligodendrocyte differentiation throughout human development, potentially influencing crucial processes such as cellular maturation and myelination in the central nervous system.

During the developmental trajectory of OPCs into MOs, transcription factors like SOX4, SOX5, SOX6, and SOX11 are active in the OPCs and downregulate during maturation. Co-expression network analysis in both infant and adult human samples (weight >0.4, <u>Supplementary file 2</u>) shows that THAP9 demonstrates an inverse expression pattern to these transcription factors. On the other hand, THAP9 is upregulated in MOs, along with important myelination markers MBP and MOG, which are expressed specifically in the late stages of oligodendrocyte differentiation. It is interesting to note that the upregulation of THAP9 along with MOG and MBP (<u>Supplementary file 3</u>) coincides with the downregulation of SOX4, SOX5, SOX6, and SOX11, as well as early specification factors and PDGFRA. This indicates that THAP9 might be involved in the regulatory network governing the maturation of oligodendrocytes or in the differentiation, maintenance and myelination process in MOs.

Most studies in oligodendrocyte biology have been carried out in mouse model systems. However, significant differences exist between human and mouse oligodendrocyte biology due to the evolutionary divergence between humans and mouse ^35^. These variations due to interspecies differences have resulted in gaps in the information, making it important to study more human-specific genes. Human and mouse oligodendrocyte biology varies in various ways, which include differences in myelination timing^36^, cell turnover rates, and molecular signatures^37^. Around two hundred genes are expressed in human oligodendrocytes but are not expressed in mouse^37^. The absence of the THAP9 gene in several rodents, including mice and rats, indicates that it may be involved in human-specific pathways related to oligodendrocyte differentiation and maturation and contribute to some of the unique features of human myelination compared to mouse models.

Our studies show that THAP9 is a promising candidate gene linked to human oligodendrocyte maturation. We also highlight the importance of species-specific genes for oligodendrocyte maturation and emphasise the importance of human-specific genes in this process. Thus, we also highlight the fundamental challenges in neurogenetic research due to the limitations of model organisms in completely understanding the molecular mechanisms involved in human neurological development.

## Methodology

### Downloading of Datasets and Preprocessing

GSE239662, GSE130063 (RNAseq) and GSE239660 (ChIP-seq) datasets from the GEO (Gene Expression Omnibus) database ^38^ were downloaded. All the raw files were downloaded from the NCBI SRA (Sequence Read Archive) using SRA-toolkit ^39^ and the prefetch (3.1.1) function. All the .sra files were converted to fastq. FastQC (version 0.12.1) ^40^ was used to control the quality of all the sequencing files. The trimming and removal of low-quality sequences of sequencing files were done using Fastp (version 0.23.4) ^41^.

The GSE239662 dataset consists of RNAseq samples obtained from the gray matter of the dorsolateral prefrontal cortex of the human brain, comprising two age groups: Adults aged 30-40 years (n=12 individuals) and Infants ages 0-2 years (n=10 individuals). Two distinct oligodendrocyte lineage populations were isolated from each individual and sequenced: Oligodendrocyte precursor cells (OPCs) and Mature oligodendrocytes (MOs). Consisting of a total of 39 RNA-sequencing files.

The GSE130063 dataset consists of RNA sequencing profiles of 24 samples (8 samples in different stages in triplicates) of induced oligodendrocyte progenitor cells (iOPCs) generated from human somatic cells. The different cell populations (A2B5+ iOPCs, O4+iOPCs, OLIG2+iOPCs, SOX10+iOPCs and Fibroblasts, Suppl. Table 1) were analysed in triplicates. A2B5(BJ): A2B5+ makers cells derived from human fibroblast (BJ) induction representing early-stage iOPCs; A2B5(DF): A2B5+ makers cells derived from human dermal fibroblasts induction representing early-stage iOPCs; OLIG2_labelled (cells with OLIG2+ maker derived from OLIG2::eGFP fibroblasts, representing completely developed OPCs), SOX10_labelled (cells with SOX10+ maker derived from SOX10::eGFP fibroblasts, representing a completely developed OPCs), O4(D40) and O4(D60), iOPCs derived from BJ on day 40 and 60; OLIG2_ESC, OLIG2::eGFP fibroblasts and DF, dermal fibroblasts. GFP+ cells had bipolar morphology. A total of 12 triplicate samples were used in the study.

The GSE239660 dataset consists of 24 ChIP-seq samples analysing H3K27ac histone modification marks in oligodendrocyte lineage populations isolated from the gray matter of the human dorsolateral prefrontal cortex. Samples were obtained from both infant and adult donors. Immunoprecipitation was performed using rabbit polyclonal anti-H3K27ac antibody to analyse the distribution of active enhancer marks. The dataset includes corresponding input controls for each experimental condition. The dataset includes samples from infants (5 MOs and 5 OPCs) and adults (5 MOs and 5 OPCs).

All the downstream analysis was carried out on R (version 4.3.1)

#### RNAseq analysis

For RNAseq analysis, transcript quantification was done using Kallisto (version 0.44.0)^42^. The abundance files were then processed in R (version 4.3.1) using tximport(version 1.28.0). Gene-level counts were obtained by summarising transcript-level abundances using “length-scaled TPM” values. Transcript-to-gene mapping was done using human genome annotations from EnsDb.Hsapiens.v86. The differential expression analysis was performed using DESeq2 (version 1.40.2) ^43^. A DESeqDataSet was constructed using raw count data with cell type as an experimental design factor. Differential expression between MOs and OPCs cell types was assessed using DESeq2 implementation of the Wald test. Genes with an adjusted p-value < 0.05 (Benjamini-Hochberg correction) were considered significantly differentially expressed.

#### Isoform analysis

The RNAseq data was processed using R version 4.3.1, and an isoform analysis was done using the IsoformSwitchAnalzeR package^44^ in R. Transcript-level quantification data was imported and analysed. The analysis was performed on the human genome assembly GRCh38 (Ensembl release 112) with corresponding transcript annotation. Differential exon usage was assessed using DEXSeq through the IsoformSwitchAnalyzeR framework.

Multiple computational tools were employed to characterise the functional potential of transcripts: Coding potential was evaluated using CPC2 (Coding Potential Calculator 2)^45^ to predict which isoforms were coding and non-coding. This tool supports Vector machine-based classifiers and uses six biologically important sequence features to make predictions. Protein domains were identified using PFam^46^, and signal peptides were predicted using SignalP-5.0 ^47^; Intrinsically disordered regions (IDRs) were identified using IUPred2A^48^, which makes sequence-based predictions of unstructured regions. Subcellular localisation prediction was performed using the DeepLoc2 ^49^ tool, which uses deep learning methods to predict cellular compartment localisation and map the cellular distribution of isoforms. Transmembrane topology was analysed using the DeepTMHMM ^50^ tool that uses deep learning architecture.

#### Gene Co-expression analysis

Weighted Gene Co-expression Network Analysis (WGCNA)^51^ (version 1.72-5) was used to characterise the transcriptional networks and identify genes showing coordinated expression patterns. The count’s matrix was normalised using Variance Stabilizing Transformation (vst). Outliers were checked using Hierarchical clustering and PCA. Soft-thresholding power (ß) was calculated to get the scale-free network. To get the appropriate power, the cut-off value for R^2^ was set to 0.8. An adjacency matrix was formed, and a Topological Overlap Matrix (TOM) was calculated using the calculated power. BlockwiseModules function was used to detect the modules, where the minimum module size was set to 30 and the maximum block size to 9000. To merge the module based on dissimilarity, the cut height was set to 0.25. Module eigengenes (MEs) were obtained and used to calculate module traits (cell type) using Pearson correlation and were displayed in the form of a heatmap. Module membership (MM) and gene significance (GS) were calculated to get more information on individual genes’ correlation with the module eigengene and traits belonging to the Turquoise module (module with the genes of interest). Genes with absolute MM and GS higher than 0.8 were selected and filtered to get the edge file. Edges with a weight of less than 0.25 were removed to get a refined network. A clusterProfiler (version 4.10.1) package was used for the functional enrichment analysis. Molecular Function (MF), Biological Process (BP), and Cellular Component (CC) were obtained for the desired module; the p-value and q-value cut-off were set to 0.05 and 0.1, respectively.

#### ChIP-seq analysis and visualization

After trimming and removing low-quality reads, all the sequence files were mapped to the human genome assembly GRCh38/hg38 using Bowtie2 (version 2.5.4)^52^. All the multi-mapping reads and low-quality alignments were removed using SAMtools (version 1.20)^53^. Read mapping to ENCODE-blacklisted genomic regions were removed using BEDTools (version 2.31.1)^54^. MACS2 (version 2.2.9.1)^55^ peak calling was used to identify H3K27-ac regions, with input controls for both cell types (OPCs and MOs) of both age groups (infants and adults). All the signal tracks were normalised to 1 million reads and converted to bigWig format (.bw files) for visualisation. The Integrative Genomics Viewer (IGV) (version 2.17.0)^56^ was used to visualise H3K27ac peaks and signal distribution across the genome to get a detailed view of chromatin modifications at specific genomic loci.

## Author Contributions

TB performed all data analyses, prepared the figures, and drafted the manuscript. DP conducted the WGCNA analysis and contributed to the interpretation of the results. As the corresponding author, SM supervised the study, provided critical revisions, and edited the manuscript. All authors reviewed and approved the final version of the manuscript.

## Code and data

No new codes were developed for this study. All tools and scripts used in the analysis have been documented and made publicly available in the [GitHub repository](https://github.com/TanujaBhardwaj/Olig_study.git).

Files attached:

Supplementary file 1

<u>Supplementary file 2</u>

<u>Supplementary file 3</u>

